# Ribosome Profiling Reveals a Dichotomy Between Ribosome Occupancy of Nuclear-Encoded and Mitochondrial-Encoded OXPHOS mRNA Transcripts in a Striatal Cell Model of Huntington Disease

**DOI:** 10.1101/2021.01.30.428960

**Authors:** S Subramaniam, N Shahani

## Abstract

Huntington disease (HD) is caused by an expanded polyglutamine mutation in huntingtin (mHTT), which promotes a prominent atrophy in the striatum and subsequent psychiatric, cognitive, and choreiform movements. Multiple lines of evidence point to an association between HD and aberrant striatal mitochondrial functions. However, present knowledge about whether (or how) mitochondrial mRNA translation is differentially regulated in HD remains unclear. We have recently applied ribosome profiling (Ribo-Seq), a technique based on the high-throughput sequencing of ribosome-protected mRNA fragments, to analyze detailed snapshots of ribosome occupancy of the mitochondrial mRNA transcripts in control and HD striatal cells. Ribo-seq data revealed almost unaltered ribosome occupancy on the nuclear-encoded mitochondrial transcripts involved in oxidative phosphorylation (OXPHOS) and only a mild reduction in ribosome occupancy on a few selected transcripts (SHDA, Ndufv1, Timm23, Tomm5, and Mrps22) in HD cells. By contrast, ribosome occupancy of mitochondrially encoded OXPHOS mRNAs (mtNd-1, mtNd-2, mtNd-4, mtNd-4l, mtNd-5, mtNd-6, mt-Co1, mtCyt b, and mt-ATP8) was dramatically increased, implying widespread dichotomous effects on ribosome occupancy and OXPHOS mRNA translation in HD. Thus, mHTT may command signals that specifically regulate translation of the mitochondrial OXPHOS transcripts and influence HD pathogenesis.

## Introduction

Expansion of the CAG repeat in the huntingtin (HTT) gene causes the motor disturbance, cognitive loss, and psychiatric manifestations of Huntington’s Disease (HD), but the exact mechanism by which mutant HTT (mHTT) induces its pathological effect in the brain remains unclear. Previous studies found widespread mitochondrial abnormalities, including decreased complex activities ^[1–8]^,, oxidative damage ^[9–10]^, mitochondrial depolarization ^[11]^, calcium defects ^[12–13]^, altered biogenesis ^[14–17]^, and mitophagy ^[18–20]^ in HD models and in patient samples. Despite several investigations, the precise cellular mode(s) of action of mHTT on mitochondria remains controversial. For example, mHTT had no effect on oxidative metabolism and mitochondrial Ca^2+^ handling in HD ^[21–22]^, whereas mHTT directly interacted with mitochondria and altered mitochondrial proteostasis ^[23–27]^. Consequently, the details of the mechanisms that relate mHTT to mitochondrial defects remain unclear.

Mitochondria are essential organelles that play a vital role in numerous cellular processes, and they contain their own semi-autonomous system of gene expression and mRNA translation machineries. The mitochondrial genome codes for components of the ribosomal-RNA genes and 22 transfer-RNA genes, as well as 13 proteins ^[28–29]^ that participate in the oxidative phosphorylation (OXPHOS) reactions of complexes (I, III, IV). Additional genes required for mtDNA maintenance, replication, transcription, translation, post-translational modification, transport, assembly, and expression of OXPHOS complexes (II, V) are exclusively encoded by the nucleus ^[30]^. Thus, mitochondrial functions, and particularly the translation and assembly of the OXPHOS complexes, are mechanistically daunting as they are encoded by different genomes. For these reasons, the molecular and biochemical control of mitochondrially encoded protein synthesis and the role of defects in this process in neurodegenerative diseases remain poorly understood.

The aim of the present study was to use our recently reported high-quality parallel RNA sequencing (RNA-seq) and ribosome profiling (Ribo-seq) technique^[31]^ to compare in vivo genome-wide information on protein synthesis (GWIPS) in healthy and HD striatal cells. Here, we report our analysis of the relative expression levels of mitochondrially encoded mRNA transcripts, ribosome density and ribosome occupancy in the OXPHOS genes.

## Results

### Ribosome profiling of ribosome-protected fragments from 55S and 80S ribosomes

We carried out systematic Ribo-Seq and RNA-seq analyses in well-established ST-Hdh-7/Q7 (WT control), ST-Hdh-Q7/Q111 (HD-het), ST-Hdh-Q111/Q111 (HD-homo) knock-in mouse striatal cell lines that express full-length wild-type (polyQ7) HTT and one (HTT-het) or two (HTT-homo) copies of mutant (polyQ111) HTT. Three replicates of WT control, HD-het, and HD-homo mutant striatal cells were subjected to ribosome profiling and the fractions contained 55S (mitoribosomes) and 80S (cytoribosomes) collected and prepared Ribo-Seq and matching RNA-Seq (Fig. 1). Through the use of multiple quality control measures, we have previously generated a high-quality global Ribo-Seq library of control, HD-het, and HD-homo cells ^[31]^.

**Figure. 1.**
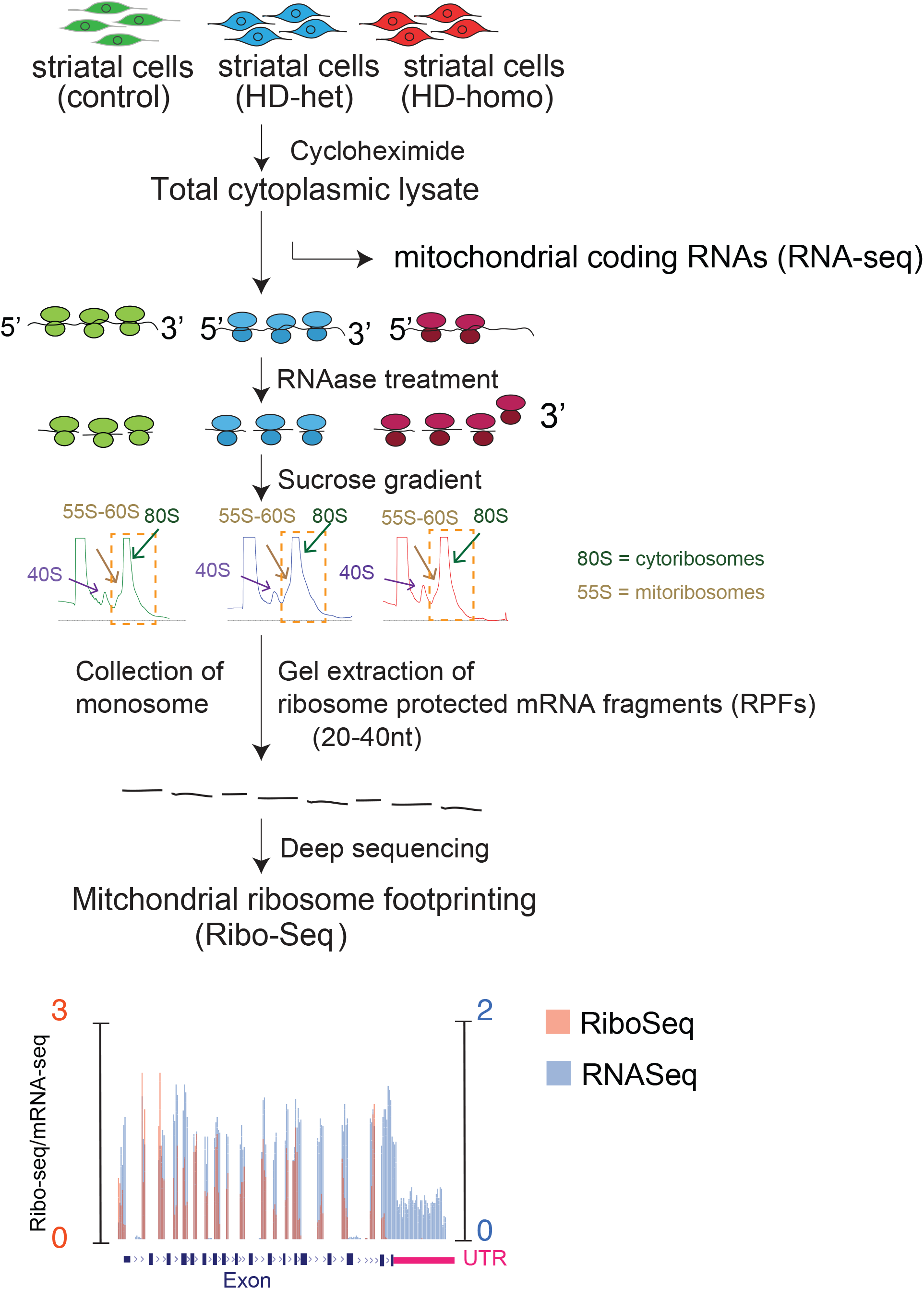
A schematic diagram showing the experimental design for performing the ribosome profiling (Ribo-Seq) and mRNA sequencing (RNA-Seq) in indicated mouse striatal cells. RNAase digested polysomes collected as 80S peaks most likely comprised of mitoribosomes with lower sedimentation coefficients (~55S, brown arrow).

Here, we explored ribosome occupancy by examining the ribosome profiles of mitochondrially encoded mRNAs in the Ribo-Seq. We combined the uploaded profiles, as a track hub in the University of California Santa Cruz (UCSC) Genome Browser 42, from the triplicate experiments and overlaid ribosome protected fragments (RPF, orange) and mRNA abundance (mRNA, blue). We then estimated the ribosome occupancy (R.O) as the ratio between CDS RPF abundance and mRNA abundance and for each gene (RPF/mRNA) from the raw read counts from the UCSC browser (Fig. 1).

### Ribosome occupancy is increased on the mitochondrially encoded OXPHOS genes in HD

We evaluated the ribosome occupancy of all 13 mitochondrially encoded OXPHOS mRNA transcripts (mt-OXPHOS transcripts). We observed ribosome occupancy for 11 mt-OXPHOS mRNA transcripts. The complex I subunits mt-Nd1, mt-Nd2, mt-Nd3, mt-Nd4, mt-Nd4l, mt-Nd5, and mt-Nd6 showed high RPF, low mRNA, and high R.O (RPF/mRNA) in HD cells compared to control; the R.O was also much higher in HD-het compared to HD-homo cells [Fig. 2(A-D), Fig. 3 (A-C)]. Mt-CyB, the only mitochondrially encoded complex III subunit, also showed high RPF and R.O (Fig. 4A). The complex IV subunit mt-Co1 and the complex V subunit mt-Atp8 showed high RFP and a significant trend of higher R.O in the HD-het and HD-homo cells (Fig. 4B, C). We were able to obtain R.O for the remaining three mt-OXPHOS transcripts (mt-Co3, mt-Co2, and mt-Atp6) and these showed a trend of higher R.O in HD cells (Supplementary Figure 1). However, we were unable to obtain statistically significant differences, as no RPF or mRNA reads were discernible in one or more replicates of the Ribo-Seq experiments (Supplementary Figure 1). Variation in the ribosome density across the transcripts was also observed; while Mt-Nd1 and mt-Nd3, mt-Nd4l showed ribosomes are preferentially located at the 5’ region mt-Nd3, mt-Nd6 and mt-CyB towards the 3’ region. These results indicate that almost all the mt-OXPHOS mRNA transcripts showed a trend of enhanced R.O with altered ribosome density in HD cells, and particularly in HD-het cells (See discussion).

**Figure 2.**
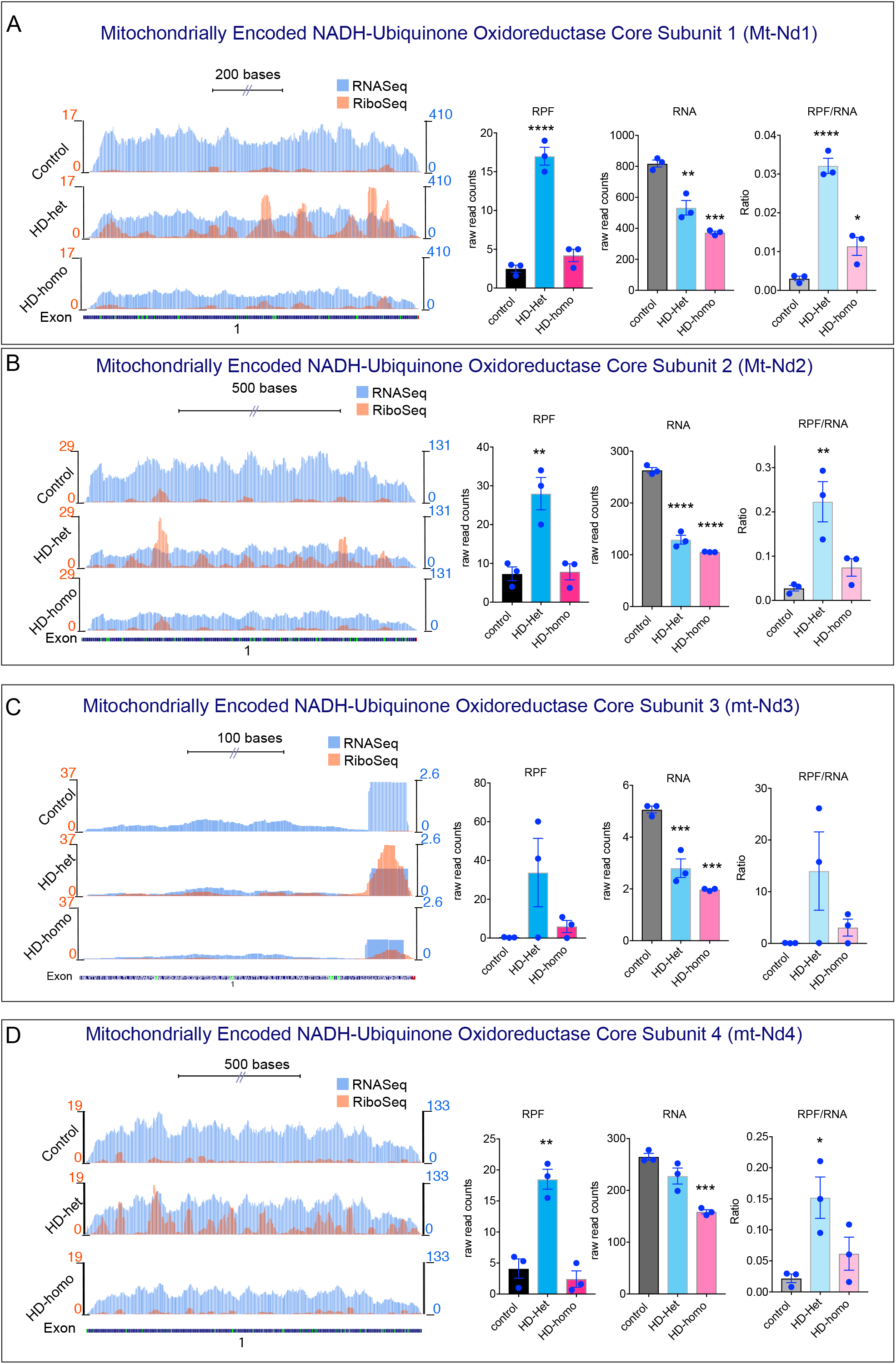
(A-D) Representative graphs showing overlay of mitochondrially-encoded transcripts Ribo-Seq (RPF)/mRNA-Seq and their corresponding average individual raw reads for RPF and mRNA and the ratio of RPF/mRNA (Ribosome occupancy) from the triple experiments for mt-ND1 (A), mt-ND2 (B), mt-ND3 (C) and mt-ND4 (D), extracted from UCSC Genome Browser. Error bar represents mean ± SEM (n = 3 independent experiments), \*\*\*\**P*< 0.0001, \*\*\**P*< 0.001, \*\**P*< 0.01, \**P*< 0.05, one-way ANOVA followed by Tukey’s multiple comparison test.

**Figure 3.**
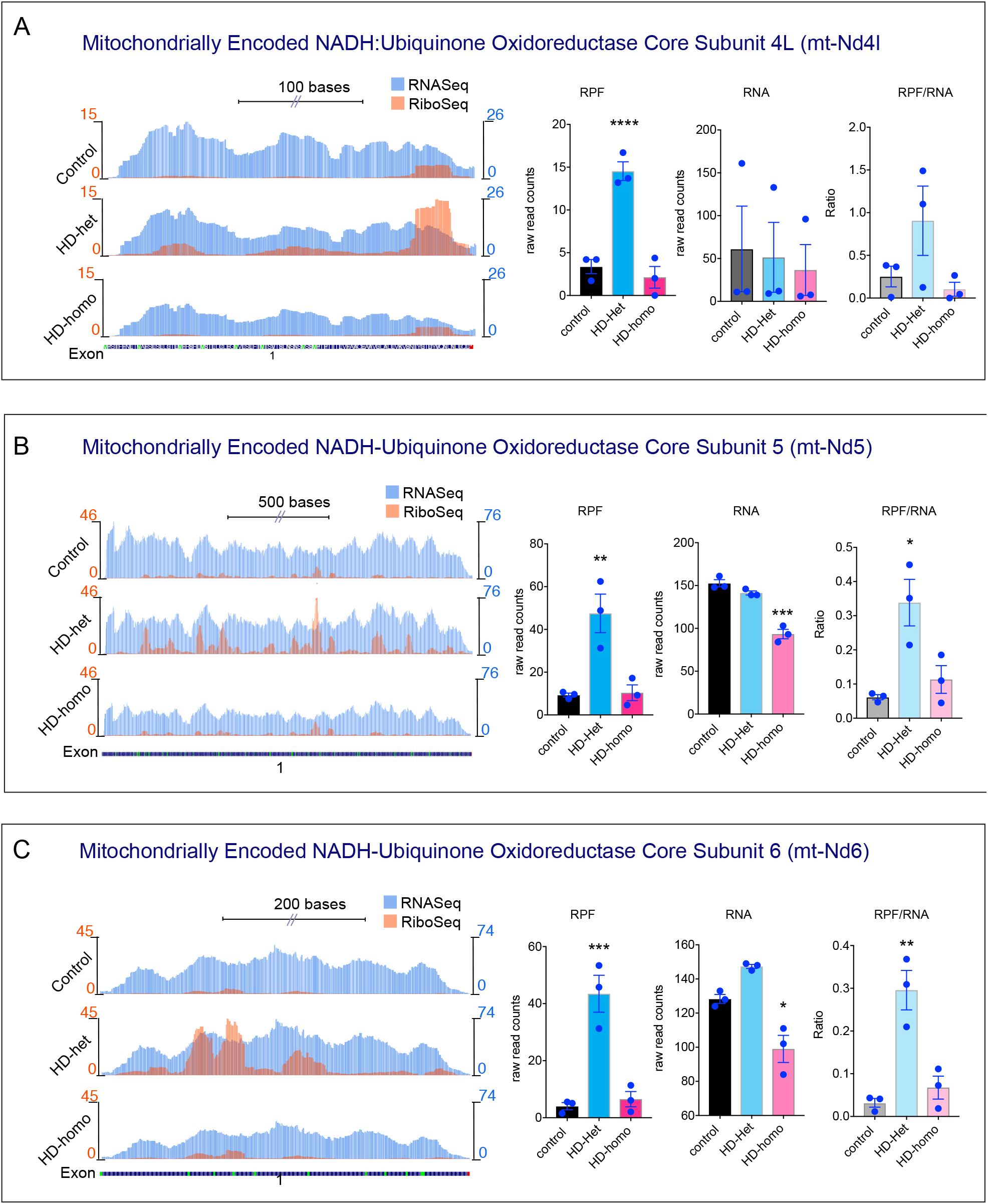
Representative graphs showing overlay of mitochondrially-encoded transcripts Ribo-Seq (RPF)/mRNA-Seq and their corresponding average individual raw reads for RPF and mRNA and the ratio of RPF/mRNA (Ribosome occupancy) from the triple experiments for mt-Nd4l (A), mt-Nd5 (B), and mt-Nd6 (C) extracted from UCSC Genome Browser. Error bar represents mean ± SEM (n = 3 independent experiments), \*\**P*< 0.01, \**P*< 0.05, one-way ANOVA followed by Tukey’s multiple comparison test.

**Figure 4.**
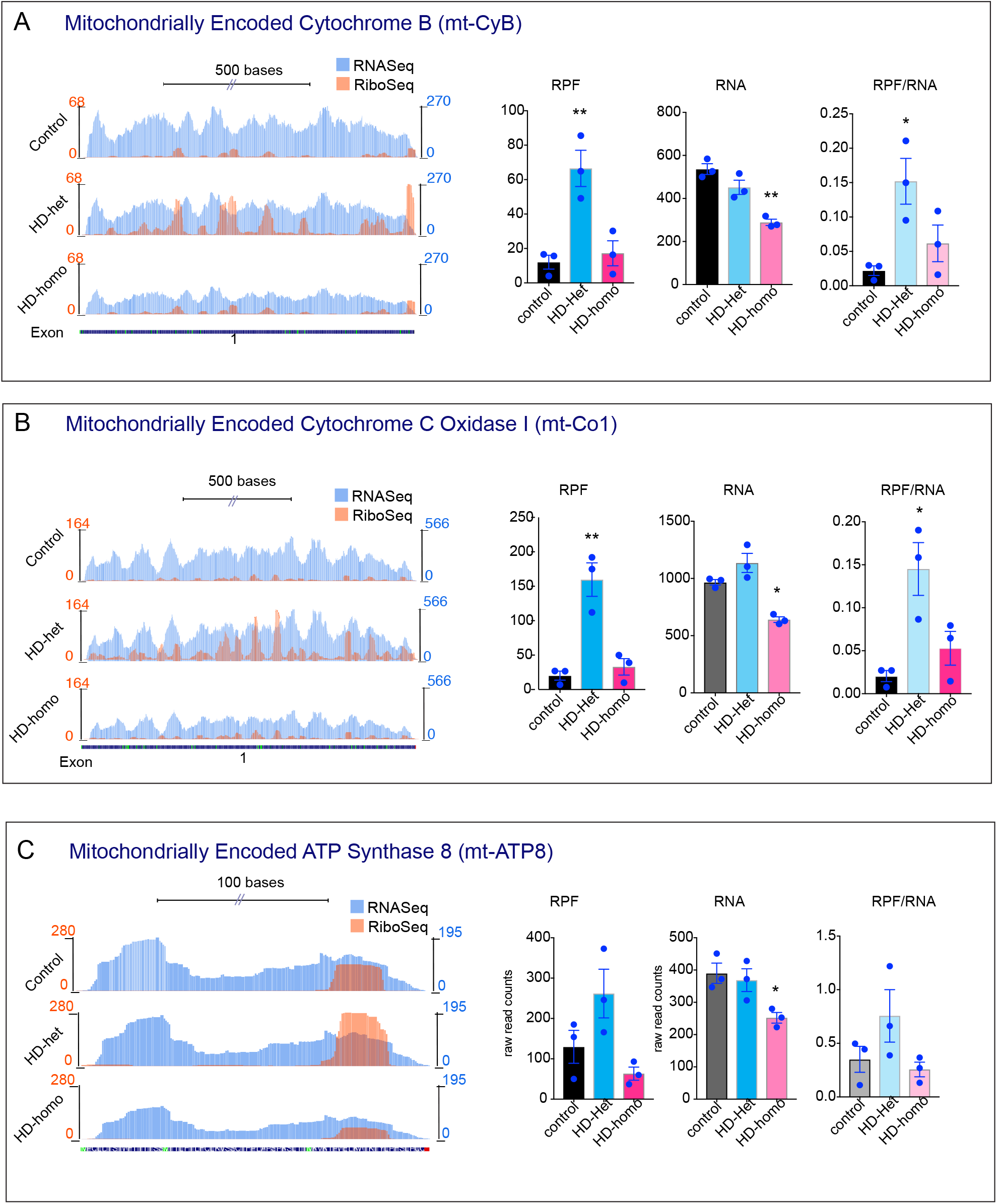
Representative graphs showing overlay of mitochondrially-encoded transcripts Ribo-Seq (RPF)/mRNA-Seq and their corresponding average individual raw reads for RPF and mRNA and the ratio of RPF/mRNA (Ribosome occupancy) from the triple experiments for mt-Co1 (A), mt-CyB (B), and mt-ATP8 (C) extracted from UCSC Genome Browser. Error bar represents mean ± SEM (n = 3 independent experiments), \*\**P*< 0.01, \**P*< 0.05, one-way ANOVA followed by Tukey’s multiple comparison test.

### Ribosome occupancy is slightly decreased or unaltered on the mitochondrially encoded OXPHOS genes in HD

We also investigated the RFP and R.O of selected nuclear-encoded complex I subunits. Ndufv1 showed ribosomes distributed throughout the 10 exons, whereas RPF is significantly diminished in HD-het cells and unaffected in HD-homo cells compared to controls (Fig. 5A). However, the R.O of the Ndufv1 transcript showed no differences (Fig. 5A). A selected example of nuclear encoded complex IV subunits (nu-Cox5a) showed no difference in the R.O between the control and HD cells (Fig. 5B). Similarly, the R.O of the complex III subunit Uqcrb was the same for control and HD cells (Fig. 5C), whereas the R.O of CytC1 was diminished in the HD-het cells compared to the controls (Fig. 6A). The Complex V subunits (ATP5o) showed no significant alterations in R.O among the cell types (Fig. 6B). For complex II, the Sdha showed a significant reduction in R.O in HD cells compared to controls (Fig. 6C), while the R.O of the Sdhc and Shhb subunits were similar between the control and HD cells (Data not shown).

**Figure 5.**
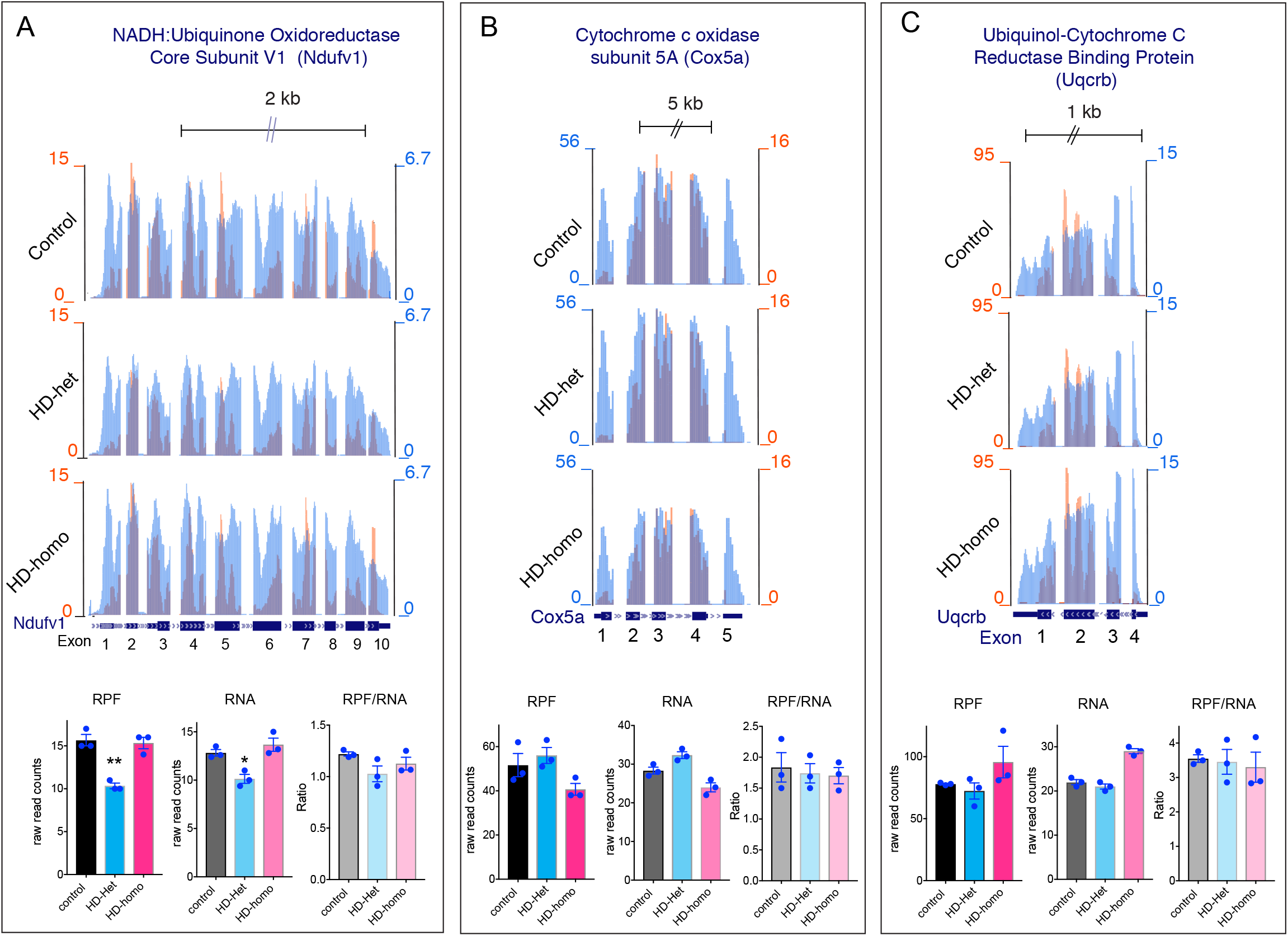
Representative graphs showing overlay of mitochondrially-encoded transcripts Ribo-Seq (RPF)/mRNA-Seq and their corresponding average individual raw reads for RPF and mRNA and the ratio of RPF/mRNA (Ribosome occupancy) from the triple experiments for Ndufv1 (A), Cox5a (B), and Uqcrb (C) extracted from UCSC Genome Browser. Error bar represents mean ± SEM (n = 3 independent experiments), \*\**P*< 0.01, \**P*< 0.05, one-way ANOVA followed by Tukey’s multiple comparison test.

**Figure 6.**
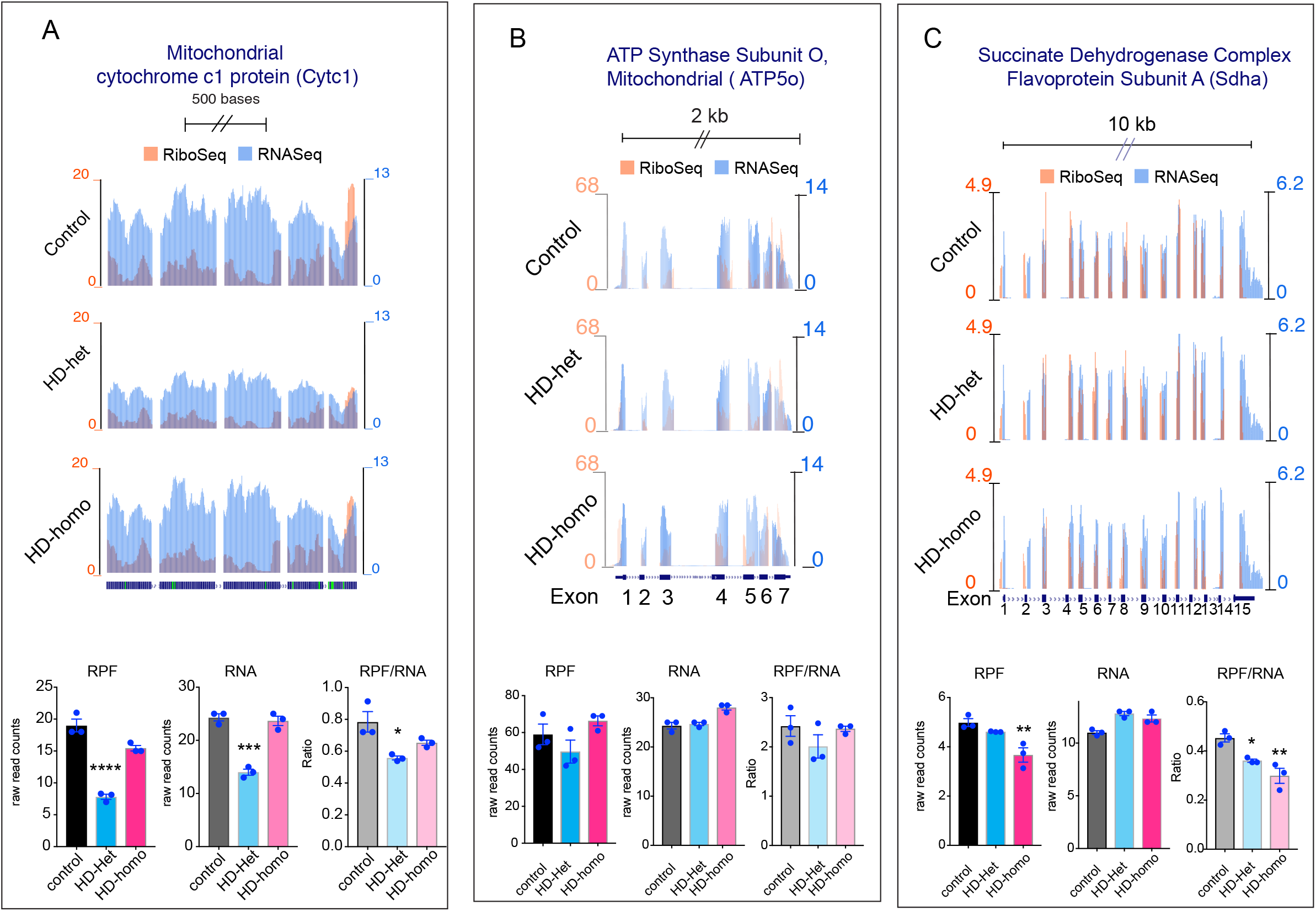
Representative graphs showing overlay of mitochondrially-encoded transcripts Ribo-Seq (RPF)/mRNA-Seq and their corresponding average individual raw reads for RPF and mRNA and the ratio of RPF/mRNA (Ribosome occupancy) from the triple experiments for Cytc1 (A), ATP5o (B), and Sdha (C) extracted from UCSC Genome Browser. Error bar represents mean ± SEM (n = 3 independent experiments), \*\*\*\**P*< 0.0001, \*\*\**P*< 0.001, \*\**P*< 0.01, \**P*< 0.05, one-way ANOVA followed by Tukey’s multiple comparison test.

### Ribosome occupancy is decreased or unaltered on the mitochondrial transcripts that code for the outer membrane, mitochondrial biogenesis, and mitochondrial ribosomes

Examination of nuclear-encoded mitochondrial outer membrane transcript Tomm5 showed significantly decreased RFA and RNA, but no significant alterations of R.O (one-way ANOVA, Tukey’s multiple comparison), indicating that the decreased occupancy was due to diminished RNA levels (Fig. 7A). Timm23 showed lower RPF in HD-het cells than in HD-homo cells but no significant changes in RNA or R.O (Fig. 7B). Interestingly, the RPF of PCK2 was increased in HD-het due to increases in mRNA, but no change was observed in R.O (Fig. 7C).

**Figure 7.**
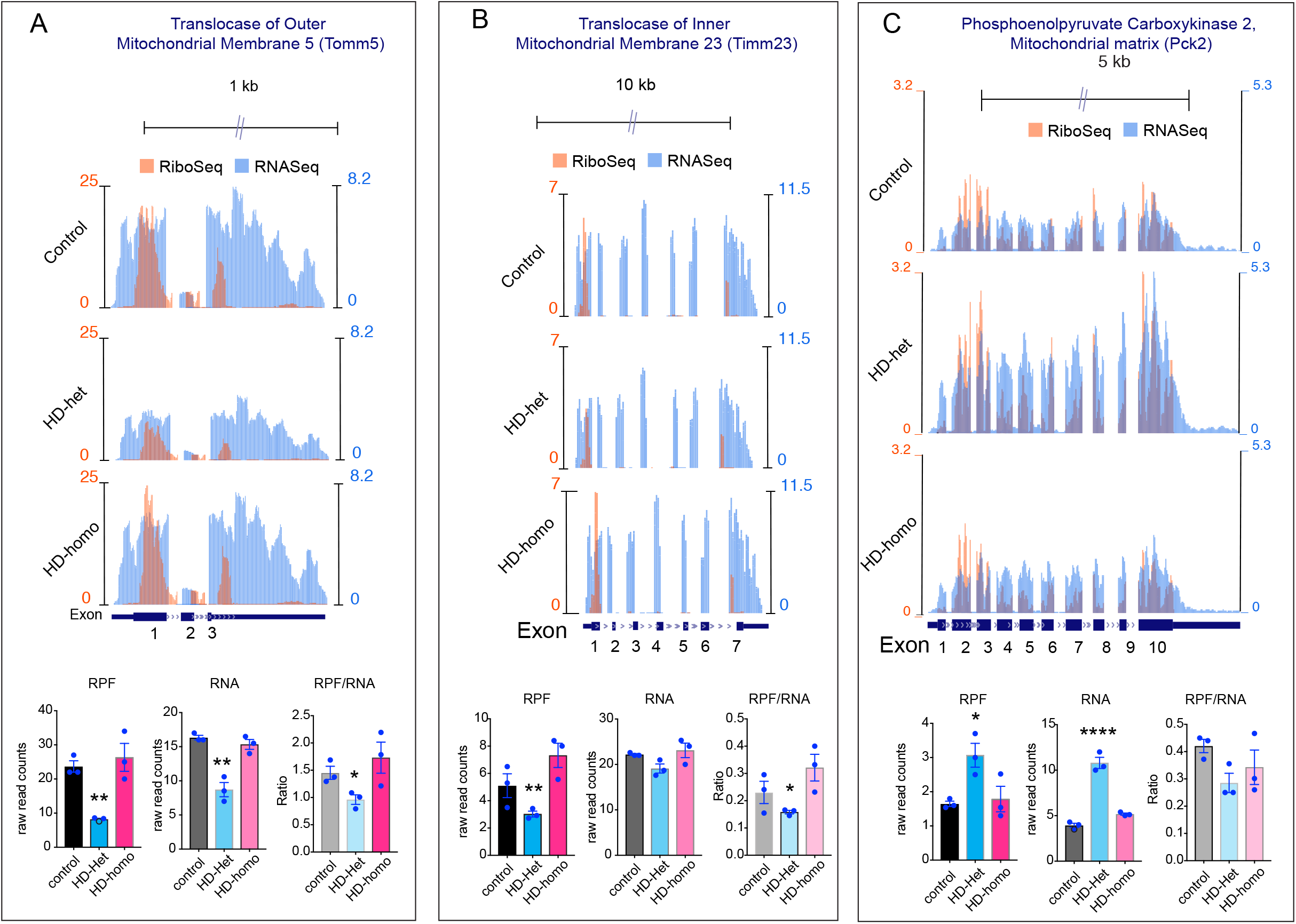
Representative graphs showing overlay of mitochondrially-encoded transcripts Ribo-Seq (RPF)/mRNA-Seq and their corresponding average individual raw reads for RPF and mRNA and the ratio of RPF/mRNA (Ribosome occupancy) from the triple experiments for Tomm5 (A), Timm23 (B), and Pck2 (C) extracted from UCSC Genome Browser. Error bar represents mean ± SEM (n = 3 independent experiments), \*\*\*\**P*< 0.0001, \*\**P*< 0.01, \**P*< 0.05, one-way ANOVA followed by Tukey’s multiple comparison test.

We also compared the RFP, RNA, and R.O of various nuclear-encoded mitochondrial transcripts, as shown in Fig. 8. The Mrps22, a mitochondrial ribosome component, showed significantly decreased RFP and R.O (Fig. 8A), while Tim14, the inner membrane translocase, showed decreased RPF and RNA but no change in R.O (Fig. 8B). No other tested nuclear-coded mitochondrial transcripts (Mrpl3, Tufm, Rars2, Opa1, Fis1, and Vdac1) showed changes in ribosome occupancy (Fig. 8C-H). Additional examples of all the nuclear-coded mitochondrial mRNA transcripts in the control, HD-het, and HD-homo cells can be found in the UCSC browser at https://genome.ucsc.edu/cgi-bin/hgTracks?hubUrl=Https://de.cyverse.org/anon-files/iplant/home/rmi2lab/Hub_Collaborations/Srini/hub.txt&genome=mm10

**Figure 8.**
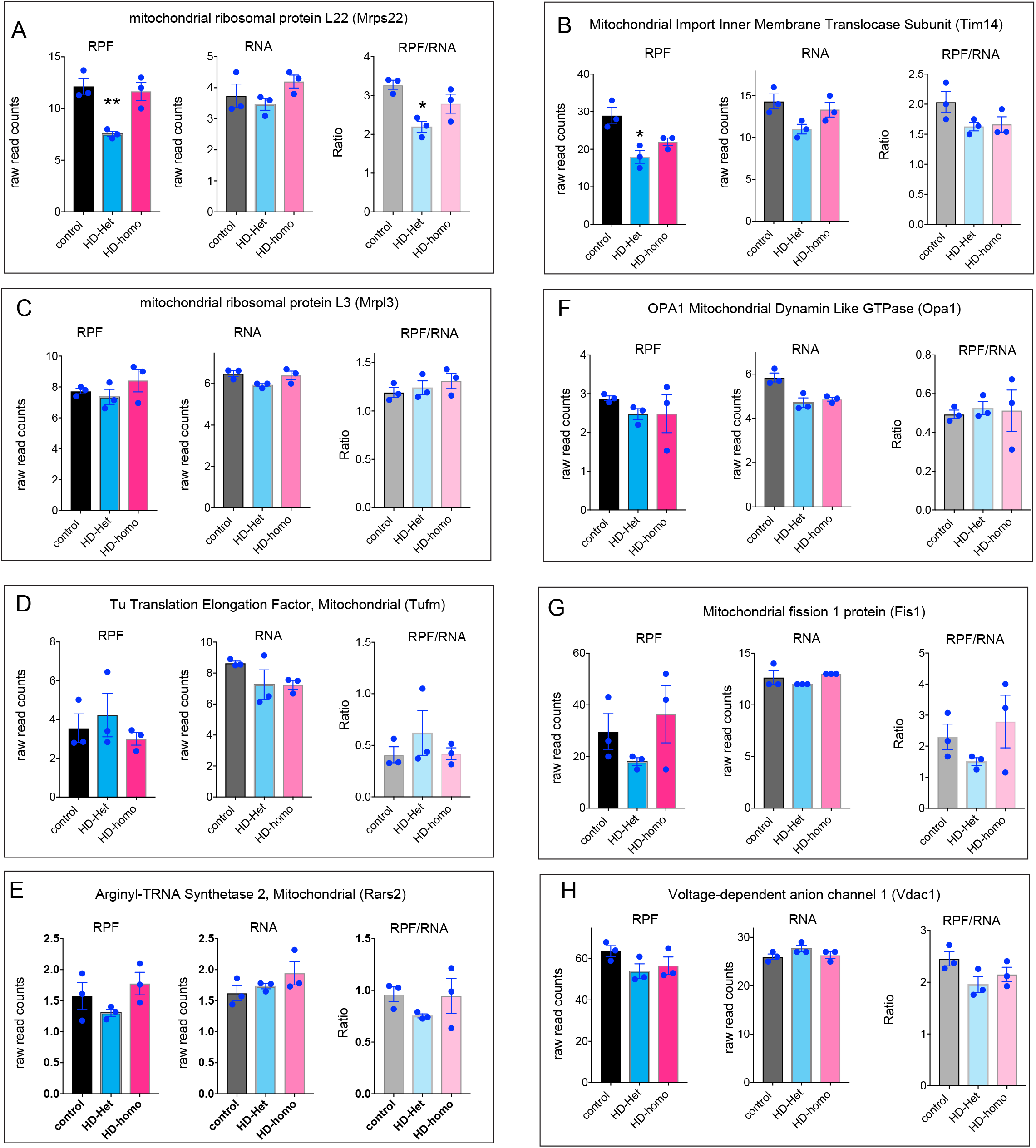
Representative individual raw reads for RPF and mRNA from the triple experiments for the indicated mRNA transcripts extracted from UCSC Genome Browser. Error bar represents mean ± SEM (n = 3 independent experiments), \*\**P*< 0.01, \**P*< 0.05, one-way ANOVA followed by Tukey’s multiple comparison test.

In summary, our ribosome profiling studies of mitochondrial transcripts in HD cells revealed an intriguing and previously unreported dichotomy: Substantial differences exist in the ribosome occupancy between nuclear-encoded vs mitochondrially encoded OXPHOS transcripts, and the ribosome occupancy levels of mitochondrially encoded OXPHOS genes is largely enhanced in HD cells. This dichotomy of ribosome occupancy implies the existence of a different mitochondrial translation mechanism in healthy versus HD cells. Understanding the possible mechanistic process behind this dichotomy mediated by mHTT (Fig. 9) can help in understanding the complex disease process and in the identification of new therapeutic targets.

**Figure 9.**
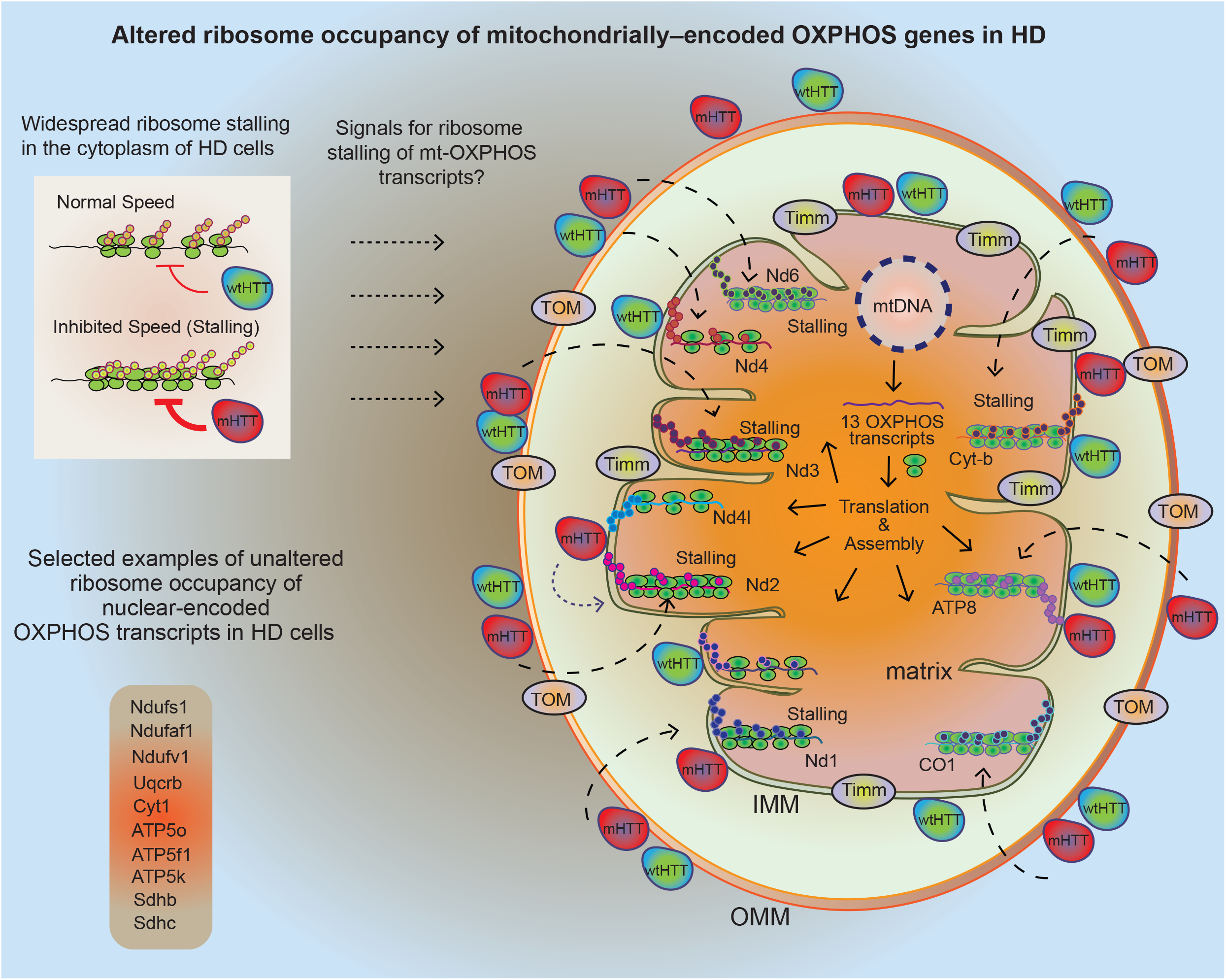
Model for mHTT-mediated ribosome stalling of mitochondrially coded OXPHOS genes. We reported widespread ribosome stalling in HD cells and our model predicted that HTT inhibits ribosome movement and mHTT gains this function and further impedes the ribosome movements and inhibits protein synthesis. Here we report the observation that an intriguing dichotomy in OXPHOS transcript ribosome loading in HD. Almost all mitochondrially encoded OXPHOS transcripts, but not the nuclear-encoded OXPHOS transcripts, show a higher ribosome occupancy in HD cells. We predict that the unique dichotomy and the associated signaling mechanisms that HTT or mHTT has on inducing mt OXPHOS transcript ribosome loading may emanant from the cytoplasm or from the inner or outer mitochondrial membrane. The possible mechanistic process for the dichotomy of ribosome occupancy in HD remains unknown.

## Discussion

Multiple aspects of mitochondrial dysfunction are linked to numerous brain diseases, and particularly neurodegenerative diseases ^[32–42]^. Perturbed mitochondrial functions and disrupted protein synthesis have been reported in HD ^[3, 16, 43–51]^, but a detailed molecular and biochemical understanding is lacking. Here, we applied ribosome profiling and obtained new insight into mitochondrial protein synthesis in HD.

We found novel mechanistic defects of translational regulation that involve enhanced loading of ribosomes onto mitochondrially coded mRNAs that control the energy production, cell signaling, and cell death functions of the mitochondria ^[52–53]^. However, this regulation was absent from the nuclear-encoded mitochondrial mRNAs, revealing an intriguing mechanistic dichotomy in the translational regulation operating between the cytoplasm and the mitochondria in the HD context.

Previous studies have indicated that both wtHTT and mHTT are localized to the mitochondria, but whether either HTT resides in the mitochondrial intermembrane space or the outer membrane remains controversial ^[18, 23, 26, 27, 54, 55]^. Nevertheless, the mitochondrial localization raises an intriguing possibility that mitochondrial membrane-associated mHTT/wtHTT can induce signaling from the inner or outer membrane surface to regulate ribosome occupancy of translating OXPHOS mRNA and the insertion of OXPHOS peptides onto the inner mitochondrial membrane. By contrast, our recent model predicted that mHTT can directly bind to the polysomes and ribosomal proteins in the cytoplasm and thereby impede the speed of ribosome translocation and protein synthesis ^[31]^. However, whether mHTT can bind to mitochondrial ribosomes (mitoribosomes) and proteins and use a similar mechanism to regulate mitochondrial mRNA translation is not clear. Nevertheless, as most (>95%) of the mitochondrial proteins are produced in the nucleus, one possibility is that mHTT may activate signals in the nucleus and cytoplasm and these then communicate messages to the mitochondrion for its own mRNA translation (Fig. 9). Recent studies in yeast showed that cytosolic translational regulators control the mitochondrial OXPHOS genes ^[56–57]^. Moreover, a novel mitoribosome-associated quality control (mtRQC) pathway comprising of mtRF-R and MTES1 is shown to rescue mitoribosomes stalling^[58]^. Further identifying the molecular nature of cellular signals and ribosome stalling mechanisms in HD cells will provide new insights that could help to curtail mitochondrial translation defects and have a potential therapeutic impact in HD.

Enhanced ribosome occupancy is predicted to increase translation efficiency and protein levels. However, this assumption is not straightforward because ribosome occupancy can also increase due to slowly elongating ribosomes and the steady-state proteins levels are modulated by posttranslational mechanisms. A previous study indicated robust differences in the mitochondrial proteome between wtHTT and mHTT cells ^[27]^, with overall proteomics diminished in the HD, consistent with our studies ^[31]^. However, they also found that the levels of selected proteins, such as mt-CO1, were increased while the levels of VDAC1 and TIM23 were diminished in mHTT-expressing cell lines. We found that the R.O of mt-CO1 is increased, the R.O of TIM23 is decreased; and the R.O of VDAC1 is unchanged. Thus, the enhanced ribosome occupancy of mRNA transcripts in HD is not directly correlated with enhanced protein levels by western blotting. This notion is further supported by the fact that the β–actin housekeeping gene shows diminished R.O and yet it shows enhanced β–actin protein levels by western blotting in HD-het cells (Supplementary Fig. 2A, B).

We also validated some of the mitochondrial proteins and found, for example, that mt-ND2 protein levels are increased (R.O is increased, Fig. 2B) in HD cells and mt-ND3 protein levels are decreased (R.O is increased, Fig. 2C), whereas mt-ND6 protein levels are unaltered (R.O is increased, Fig. 3C) (Supplementary Fig. 2B). The lack of a direct correlation between ribosome occupancy and protein levels indicates that the ribosome loading on mRNA transcripts and the steady-state-levels of fully translated proteins in HD are presumably controlled by independent mechanisms.

One important observation from this study is the differential influence of ribosome occupancy on mitochondrially coded mRNA by one copy of mHTT (HD-het) versus two copies of mHTT (HD-homo). The HD-het cells display a significantly higher ribosome occupancy on the mt-OXPHOS genes, with a minority of nu-coded OXPHOS transcripts exhibiting diminished occupancy compared to HD-homo cells (Figs. 2–4). This observation suggests that the combined presence of wtHTT and mHTT may lead to the emergence of an additive risk for translation regulatory phenotypes that readapt to the ongoing HD-related cellular demands. In addition, as wtHTT plays a key role as a negative regulator of ribosome translocation ^[31]^, this raises the possibility that enhanced ribosome occupancy in HD-het may be involved in the subtle loss of normal HTT function due to mHTT.

Alternatively, mHTT may exert a gain of function, so that the presence of two copies of the mHTT gene worsens the degree of ribosome occupancy at mt-coded RNAs and leads to stalling of ribosomes and possibly the eventual degradation of the stalled transcripts. This notion is supported by the data showing significantly reduced amounts of mitochondrial mRNA transcripts in the HD-homo versus the HD-het or control cells (Figs. 2–4). Future studies should investigate the mechanisms that may induce a decay of ribosome-stalled mitochondrial transcripts and associated factors in HD.

Taken together, our study findings demonstrate an unexpected dichotomy of ribosome occupancy among nuclear and mitochondrially coded OXPHOS transcripts due to the presence of mHTT. Defining the underlying mechanisms that create this dichotomy of OXPHOS mRNA translation and their influence on mitochondrial structure and function will reveal new understanding of HD pathogenesis and identify new therapeutic targets.

## Material and methods

### Cell culture

Mouse striatal cells (ST*Hdh*) expressing knock-in wild-type Htt^exon1^ with 7 Glu repeats (control; ST*Hdh*^*Q7/Q7*^) or expressing knock-in mutant human Htt^exon1^ with 111 Glu repeats (HD-het; ST*Hdh*^*Q7/Q111*^, and HD-homo; ST*Hdh*^*Q111/Q111*^) ^[59]^ were purchased from Coriell Institute for Medical Research (Camden, New Jersey, USA) and cultured in 10% FBS, DMEM, high glucose, 5% CO_2_, at 33°C, as described before ^[60]^.

### Isolation of ribosomes (mitoribosomes and cytoribosomes) for profiling

Global RNase foot-printings were performed during three independent rounds of cell cultures (n=3). For each round of global foot printing, mouse immortalized striatal cells (i.e. control, HD-het, and HD-homo cells) were plated in 15 cm dishes at a confluency of 70%. The following day the mediums were changed and after 2 hours the cells were incubated with cycloheximide (CHX, 100 μg/ml) for 10 min as in previous studies^[31, 61, 62]^. Cells were then scraped and washed with cold PBS (containing 100 μg/ml CHX) twice. During the second wash 5% of cells were transferred to new tubes and were lyzed by adding 700 μl of QIAzol lysis reagent. Total RNAs of these samples were isolated using miRNeasy Mini Kit (Qiagen) for total mRNA sequencing. After the second wash, the rest of the cells were lyzed in a lysis buffer containing 20 mM HEPES pH 7.3, 150 mM KCl, 10 mM MgCl_2_, 2 mM DTT, 100 μg/ml CHX, 0.5% v/v Triton X-100, 20 U/ml RNasin and EDTA free protease inhibitor cocktail (Roche). The cell lysates were passed 20 times through a 26G needle and incubated on ice for 15 minutes, then centrifuged at 21000 rpm for 15 minutes. Supernatants were transferred to new tubes. Equal total RNA amount of each sample was used for global RNase foot printing as follow; for each A260 absorbance unit of the lysates 60 units of RNaseT1 (ThermoFisher Scientific) and 0.6 μl of RNaseA (Ambion) were added and the samples were incubated at 25°C for 30 min. RNase treated samples were immediately loaded on 10-50% sucrose gradients and centrifuged at 40000 RPM (SW41Ti rotor) at 4°C for 2 hours. Gradients were fractionated using a gradient fractionator and UA-6 detector, 254 nm filter (ISCO/BRANDEL). Fractions containing both 55S (mitoribosomes) and 80S (cytoribosomes) peaks of each sample were collected and their RNAs were isolated using a miRNeasy Mini Kit (Qiagen).

### Generation of cDNA libraries from ribosome protected mRNAs

The following procedure were performed for all the RNA samples simultaneously. 20 μg of each sample was run on a 15% TBE-Urea gel (Novex) along with 26 and 32 nt RNA markers. The gel containing each sample was excised between two markers. RNAs were extracted from gel pieces by incubating gel slurries with nuclease-free water overnight at 4°C and precipitated using RNase-free isopropanol and then eluted in nuclease-free water. T4 Polynucleotide Kinase (NEB) was used to catalyze the addition of 5’ monophosphate and removal of the 3’ phosphate in the RNA fragments to leave a 3’ hydroxyl terminal needed for adapter ligation. RNA was purified using the Zymo clean and conc-5 kit (Zymo Research, Cat. # R1013). Ribosomal RNA was depleted from the samples using TruSeq total RNA rRNA-depletion protocol (Illumina, Cat. #RS-122-2201) and then RNA samples were purified using Agencourt RNAClean XP beads (Beckman Coulter).

### Generation of cDNA libraries and sequencing

NEXTflex small RNA-seq Kit v3 (Perkin Elmer) was used to ligate 5’ and 3’ adapters to purified RPF fragments, which then were reverse transcribed and amplified (14 cycles) to generate cDNA libraries. Libraries were cleaned up using NEXTflex Cleanup beads, pooled and sequenced in the NextSeq 500 (V2) using single-end 50bp chemistry at the Scripps Genomic Core, at Florida, USA.

### Generation of mRNA-seq libraries

NEBNext Ultra II Directional kit (NEB, Cat. # E776) with the NEBNext poly(A) mRNA Magnetic isolation module (NEB, Cat. # E7490) was used generate mRNA-seq libraries. Briefly, 400ng of high-quality total RNA was used to purify poly(A) mRNA, fragmented, reverse-transcribed with random primers, adapter ligated, and amplified according to manufacturer recommendations. The final libraries were validated on the bioanalyzer, pooled, and sequenced on the NextSeq 500 using paired-end 40bp chemistry.

### Ribo-Seq, RNA-seq quality control and mapping the reads to UCSC browser

RNAseq reads were trimmed using Cutadapt^[63]^ with the following parameters : -a AGATCGGAAGAGCACACGTCTGAACTCCAGTCA -A AGATCGGAAGAGCGTCGTGTAGGGAAAGAGTGT --minimum-length=15 -pair-filter=any. For Riboseq reads, 3’ adapters were trimmed using Cutadapt with the following parameters : -a TGGAATTCTCGGGTGCCAAGG --minimum-length 23. The reads were further trimmed using Cutadapt to remove 4 bases from either side of each read accordingly to the NEXTflex™ Small RNA Trimming Instructions (cutadapt -u 4 -u -4). Fastq files were checked for quality control with FastQC. Both RNAseq and Riboseq reads were next mapped to a library of mouse rRNA and tRNA sequences using Bowtie v1.1.2. Any reads mapping to these abundant contaminants were filtered out. Remaining reads were then aligned to the mouse transcriptome with RSEM v1.3.0^[64]^ using the GRCm38.p5 genome annotation and the comprehensive gene annotation from Gencode (M16 release) as transcriptome reference. Reads with a mapping quality <5 were discarded. Cleaned bam files were converted to bigWig files with Bedtools^[65]^ for visualisation using the UCSC Genome Browser. For the euclidian distance analyses, gene expression was quantified with RSEM and comparison plots were generated in R using DESeq2^[66]^ and ggplot2 packages.

### Western blot analysis

The cells were lyzed in the lysis buffer containing 20 mM HEPES pH 7.3, 150 mM KCl, 10 mM MgCl_2_, 2 mM DTT, 100 μg/ml CHX, 0.5% v/v Triton X-100, 20 U/ml RNasin and EDTA free protease inhibitor cocktail (Roche) and an RNA concentration A260 reading of 10 OD, loaded on a 30-50% sucrose gradient. Individual fractions (250 μl) were collected, the protein was precipitated using methanol/chloroform method, and loaded for Western blots analysis using antibodies to detect indicated endogenous protein. The antibodies against Actin (sc47778, 1:20000) was from Santa Cruz and against mitochondrial proteins, Mt-CO2 (A11154, 1:1000), Mt-ND2 (A6180, 1:1000), Mt-ND3 (A9940, 1:1000), Mt-ND6 (A17991, 1:1000) were from ABclonal.

### Statistical analysis

Data were expressed as mean ± SEM as indicated. Experiments were performed in biological triplicates. Statistical analysis was performed with a Student’s *t-*test or one-way ANOVA followed by Tukey’s multiple comparison test.

## Author contributions

S.S made the initial observations and conceptualized the project. N.S contributed to the Western blotting. S.S analyzed the data and wrote the paper with input from N.S.

## Data and materials availability

All the data is available in the main text or the supplementary materials.

---

**Supplementary Figure 1.**
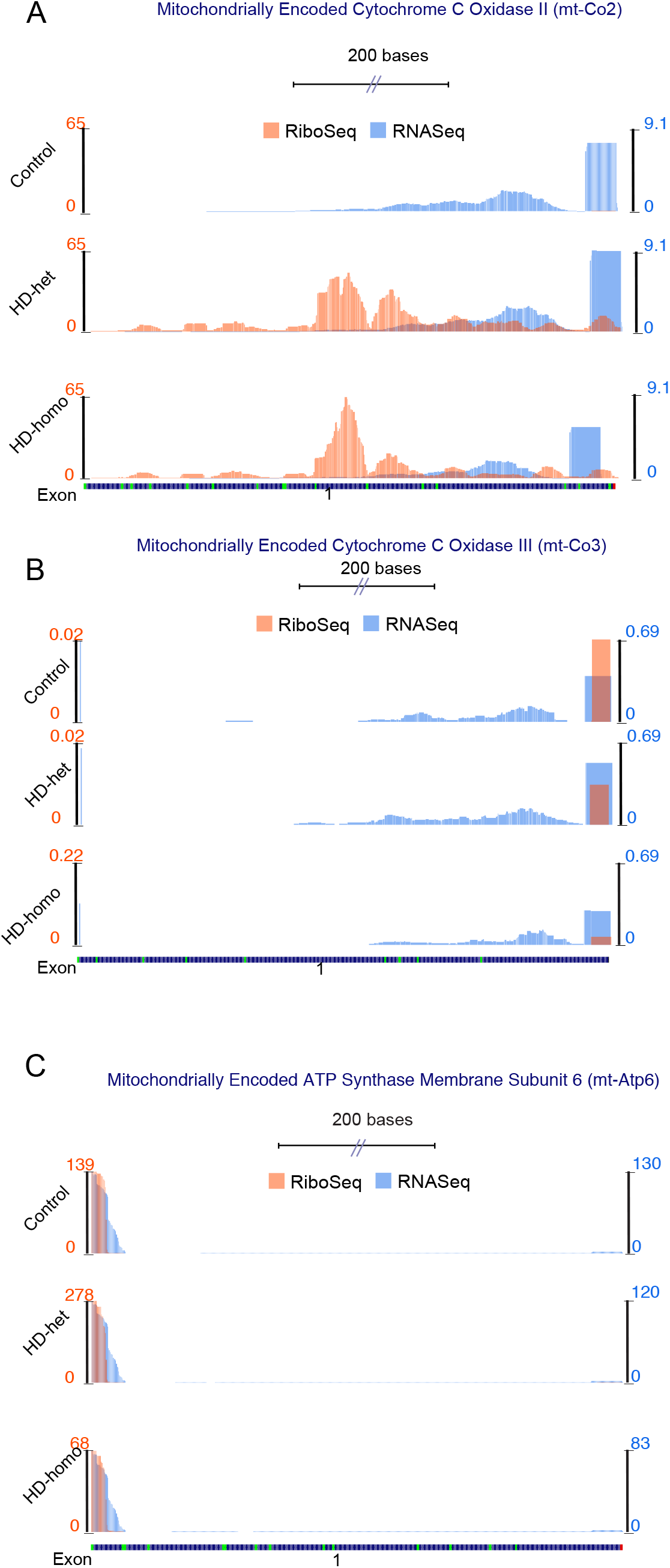
Representative graphs showing overlay of mitochondrially-encoded transcripts Ribo-Seq (RPF)/mRNA-Seq and their corresponding average individual raw reads for RPF and mRNA and the ratio of RPF/mRNA (Ribosome occupancy) from the triple experiments for mt-Co2 (A), mt-Co3 (B), and mt-Atp6 (C) extracted from UCSC Genome Browser.

**Supplementary Figure 2.**
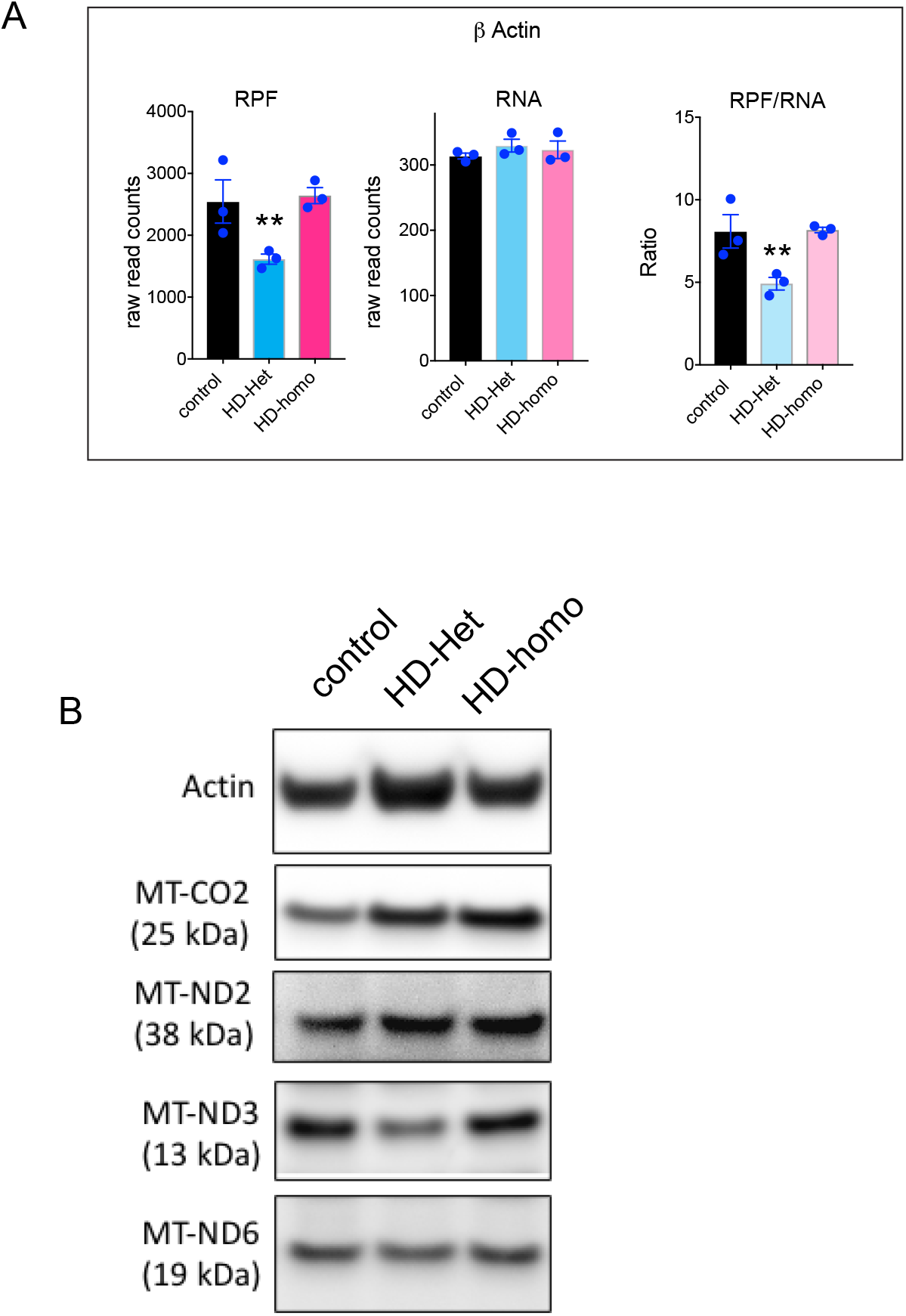
**A.** β-actin raw reads for RPF and mRNA from the triple experiments extracted from UCSC Genome Browser. Error bar represents mean ± SEM (n = 3 independent experiments), \*\**P*< 0.01, one-way ANOVA followed by Tukey’s multiple comparison test. **B**. Representative Western blot for the indicated proteins from control, HD-het and HD-homo cells.

## References

Sorbi S, Bird ED, Blass JP. Decreased pyruvate dehydrogenase complex activity in Huntington and Alzheimer brain. Ann Neurol. 1983;13(1):72–8. Epub 1983/01/01. doi: 10.1002/ana.410130116. PubMed PMID: 6219611.

Parker WD, Jr., Boyson SJ, Luder AS, Parks JK. Evidence for a defect in NADH: ubiquinone oxidoreductase (complex I) in Huntington’s disease. Neurology. 1990;40(8):1231–4. Epub 1990/08/01. doi: 10.1212/wnl.40.8.1231. PubMed PMID: 2143271.

Gu M, Gash MT, Mann VM, Javoy-Agid F, Cooper JM, Schapira AH. Mitochondrial defect in Huntington’s disease caudate nucleus. Ann Neurol. 1996;39(3):385–9. Epub 1996/03/01. doi: 10.1002/ana.410390317. PubMed PMID: 8602759.

Arenas J, Campos Y, Ribacoba R, Martin MA, Rubio JC, Ablanedo P, Cabello A. Complex I defect in muscle from patients with Huntington’s disease. Ann Neurol. 1998;43(3):397–400. Epub 1998/03/20. doi: 10.1002/ana.410430321. PubMed PMID: 9506560.

Aidt FH, Nielsen SM, Kanters J, Pesta D, Nielsen TT, Norremolle A, Hasholt L, Christiansen M, Hagen CM. Dysfunctional mitochondrial respiration in the striatum of the Huntington’s disease transgenic R6/2 mouse model. PLoS Curr. 2013;5. Epub 2013/04/10. doi: 10.1371/currents.hd.d8917b4862929772c5a2f2a34ef1c201. PubMed PMID: 23568011; PMCID: PMC3614423.

Damiano M, Diguet E, Malgorn C, D’Aurelio M, Galvan L, Petit F, Benhaim L, Guillermier M, Houitte D, Dufour N, Hantraye P, Canals JM, Alberch J, Delzescaux T, Deglon N, Beal MF, Brouillet E. A role of mitochondrial complex II defects in genetic models of Huntington’s disease expressing N-terminal fragments of mutant huntingtin. Hum Mol Genet. 2013;22(19):3869–82. Epub 2013/05/31. doi: 10.1093/hmg/ddt242. PubMed PMID: 23720495; PMCID: PMC3766181.

Pandey M, Mohanakumar KP, Usha R. Mitochondrial functional alterations in relation to pathophysiology of Huntington’s disease. J Bioenerg Biomembr. 2010;42(3):217–26. Epub 2010/05/14. doi: 10.1007/s10863-010-9288-5. PubMed PMID: 20464463.

Weydt P, Pineda VV, Torrence AE, Libby RT, Satterfield TF, Lazarowski ER, Gilbert ML, Morton GJ, Bammler TK, Strand AD, Cui L, Beyer RP, Easley CN, Smith AC, Krainc D, Luquet S, Sweet IR, Schwartz MW, La Spada AR. Thermoregulatory and metabolic defects in Huntington’s disease transgenic mice implicate PGC-1alpha in Huntington’s disease neurodegeneration. Cell Metab. 2006;4(5):349–62. Epub 2006/10/24. doi: 10.1016/j.cmet.2006.10.004. PubMed PMID: 17055784.

Polidori MC, Mecocci P, Browne SE, Senin U, Beal MF. Oxidative damage to mitochondrial DNA in Huntington’s disease parietal cortex. Neurosci Lett. 1999;272(1):53–6. Epub 1999/10/03. doi: 10.1016/s0304-3940(99)00578-9. PubMed PMID: 10507541.

Zheng J, Winderickx J, Franssens V, Liu B. A Mitochondria-Associated Oxidative Stress Perspective on Huntington’s Disease. Front Mol Neurosci. 2018;11:329. Epub 2018/10/05. doi: 10.3389/fnmol.2018.00329. PubMed PMID: 30283298; PMCID: PMC6156126.

Sawa A, Wiegand GW, Cooper J, Margolis RL, Sharp AH, Lawler JF, Jr., Greenamyre JT, Snyder SH, Ross CA. Increased apoptosis of Huntington disease lymphoblasts associated with repeat length-dependent mitochondrial depolarization. Nat Med. 1999;5(10):1194–8. Epub 1999/09/30. doi: 10.1038/13518. PubMed PMID: 10502825.

Panov AV, Gutekunst CA, Leavitt BR, Hayden MR, Burke JR, Strittmatter WJ, Greenamyre JT. Early mitochondrial calcium defects in Huntington’s disease are a direct effect of polyglutamines. Nat Neurosci. 2002;5(8):731–6. Epub 2002/06/29. doi: 10.1038/nn884. PubMed PMID: 12089530.

Bezprozvanny I, Hayden MR. Deranged neuronal calcium signaling and Huntington disease. Biochem Biophys Res Commun. 2004;322(4):1310–7. Epub 2004/09/01. doi: 10.1016/j.bbrc.2004.08.035. PubMed PMID: 15336977.

Shirendeb UP, Calkins MJ, Manczak M, Anekonda V, Dufour B, McBride JL, Mao P, Reddy PH. Mutant huntingtin’s interaction with mitochondrial protein Drp1 impairs mitochondrial biogenesis and causes defective axonal transport and synaptic degeneration in Huntington’s disease. Hum Mol Genet. 2012;21(2):406–20. Epub 2011/10/15. doi: 10.1093/hmg/ddr475. PubMed PMID: 21997870; PMCID: PMC3276281.

Haun F, Nakamura T, Shiu AD, Cho DH, Tsunemi T, Holland EA, La Spada AR, Lipton SA. S-nitrosylation of dynamin-related protein 1 mediates mutant huntingtin-induced mitochondrial fragmentation and neuronal injury in Huntington’s disease. Antioxid Redox Signal. 2013;19(11):1173–84. Epub 2013/05/07. doi: 10.1089/ars.2012.4928. PubMed PMID: 23641925; PMCID: PMC3785802.

Kim J, Moody JP, Edgerly CK, Bordiuk OL, Cormier K, Smith K, Beal MF, Ferrante RJ. Mitochondrial loss, dysfunction and altered dynamics in Huntington’s disease. Hum Mol Genet. 2010;19(20):3919–35. Epub 2010/07/28. doi: 10.1093/hmg/ddq306. PubMed PMID: 20660112; PMCID: PMC2947400.

Guo X, Disatnik MH, Monbureau M, Shamloo M, Mochly-Rosen D, Qi X. Inhibition of mitochondrial fragmentation diminishes Huntington’s disease-associated neurodegeneration. J Clin Invest. 2013;123(12):5371–88. Epub 2013/11/16. doi: 10.1172/JCI70911. PubMed PMID: 24231356; PMCID: PMC3859413.

Guo X, Sun X, Hu D, Wang YJ, Fujioka H, Vyas R, Chakrapani S, Joshi AU, Luo Y, Mochly-Rosen D, Qi X. VCP recruitment to mitochondria causes mitophagy impairment and neurodegeneration in models of Huntington’s disease. Nat Commun. 2016;7:12646. Epub 2016/08/27. doi: 10.1038/ncomms12646. PubMed PMID: 27561680; PMCID: PMC5007466 filed. The authors declare no conflict of interest.

Khalil B, El Fissi N, Aouane A, Cabirol-Pol MJ, Rival T, Lievens JC. PINK1-induced mitophagy promotes neuroprotection in Huntington’s disease. Cell Death Dis. 2015;6:e1617. Epub 2015/01/23. doi: 10.1038/cddis.2014.581. PubMed PMID: 25611391; PMCID: PMC4669776.

Hwang S, Disatnik MH, Mochly-Rosen D. Impaired GAPDH-induced mitophagy contributes to the pathology of Huntington’s disease. EMBO Mol Med. 2015;7(10):1307–26. Epub 2015/08/14. doi: 10.15252/emmm.201505256. PubMed PMID: 26268247; PMCID: PMC4604685.

Hamilton J, Brustovetsky T, Sridhar A, Pan Y, Cummins TR, Meyer JS, Brustovetsky N. Energy Metabolism and Mitochondrial Superoxide Anion Production in Pre-symptomatic Striatal Neurons Derived from Human-Induced Pluripotent Stem Cells Expressing Mutant Huntingtin. Mol Neurobiol. 2020;57(2):668–84. Epub 2019/08/23. doi: 10.1007/s12035-019-01734-2. PubMed PMID: 31435904.

Hamilton J, Brustovetsky T, Brustovetsky N. Mutant huntingtin fails to directly impair brain mitochondria. J Neurochem. 2019;151(6):716–31. Epub 2019/08/17. doi: 10.1111/jnc.14852. PubMed PMID: 31418857; PMCID: PMC6917837.

Orr AL, Li S, Wang CE, Li H, Wang J, Rong J, Xu X, Mastroberardino PG, Greenamyre JT, Li XJ. N-terminal mutant huntingtin associates with mitochondria and impairs mitochondrial trafficking. J Neurosci. 2008;28(11):2783–92. Epub 2008/03/14. doi: 10.1523/JNEUROSCI.0106-08.2008. PubMed PMID: 18337408; PMCID: PMC2652473.

Ratovitski T, Chighladze E, Arbez N, Boronina T, Herbrich S, Cole RN, Ross CA. Huntingtin protein interactions altered by polyglutamine expansion as determined by quantitative proteomic analysis. Cell Cycle. 2012;11(10):2006–21. Epub 2012/05/15. doi: 10.4161/cc.20423. PubMed PMID: 22580459; PMCID: PMC3359124.

Wang X, Zhu S, Pei Z, Drozda M, Stavrovskaya IG, Del Signore SJ, Cormier K, Shimony EM, Wang H, Ferrante RJ, Kristal BS, Friedlander RM. Inhibitors of cytochrome c release with therapeutic potential for Huntington’s disease. J Neurosci. 2008;28(38):9473–85. Epub 2008/09/19. doi: 10.1523/JNEUROSCI.1867-08.2008. PubMed PMID: 18799679; PMCID: PMC2632939.

Choo YS, Johnson GV, MacDonald M, Detloff PJ, Lesort M. Mutant huntingtin directly increases susceptibility of mitochondria to the calcium-induced permeability transition and cytochrome c release. Hum Mol Genet. 2004;13(14):1407–20. Epub 2004/05/28. doi: 10.1093/hmg/ddh162. PubMed PMID: 15163634.

Yablonska S, Ganesan V, Ferrando LM, Kim J, Pyzel A, Baranova OV, Khattar NK, Larkin TM, Baranov SV, Chen N, Strohlein CE, Stevens DA, Wang X, Chang YF, Schurdak ME, Carlisle DL, Minden JS, Friedlander RM. Mutant huntingtin disrupts mitochondrial proteostasis by interacting with TIM23. Proc Natl Acad Sci U S A. 2019;116(33):16593–602. Epub 2019/07/28. doi: 10.1073/pnas.1904101116. PubMed PMID: 31346086; PMCID: PMC6697818.

Fisher KE, Bradbury SP, Coates BS. Prediction of mitochondrial genome-wide variation through sequencing of mitochondrion-enriched extracts. Sci Rep. 2020;10(1):19123. Epub 2020/11/07. doi: 10.1038/s41598-020-76088-0. PubMed PMID: 33154458; PMCID: PMC7645498.

Wolstenholme DR. Genetic novelties in mitochondrial genomes of multicellular animals. Curr Opin Genet Dev. 1992;2(6):918–25. Epub 1992/12/01. doi: 10.1016/s0959-437x(05)80116-9. PubMed PMID: 1282405.

Reinecke F, Smeitink JA, van der Westhuizen FH. OXPHOS gene expression and control in mitochondrial disorders. Biochim Biophys Acta. 2009;1792(12):1113–21. Epub 2009/04/25. doi: 10.1016/j.bbadis.2009.04.003. PubMed PMID: 19389473.

Eshraghi M, Karunadharma P, Blin J, Shahani N, Ricci EP, Michel A, Urban N, Galli N, Sharma M, Ramírez-Jarquín UN, Florescu K, Hernandez J, and Subramaniam S. Mutant Huntingtin Stalls Ribosomes and Represses Protein Synthesis in a Cellular Model of Huntington disease. Nat Communication. 2020;In Press, BioRxiv. doi: https://doi.org/10.1101/629667.

Rothman SM. Mutations of the mitochondrial genome: clinical overview and possible pathophysiology of cell damage. Biochem Soc Symp. 1999;66:111–22. Epub 2000/09/16. doi: 10.1042/bss0660111. PubMed PMID: 10989662.

Krieger C, Duchen MR. Mitochondria, Ca2+ and neurodegenerative disease. Eur J Pharmacol. 2002;447(2-3):177–88. Epub 2002/08/02. doi: 10.1016/s0014-2999(02)01842-3. PubMed PMID: 12151010.

Swerdlow RH, Khan SM. A “mitochondrial cascade hypothesis” for sporadic Alzheimer’s disease. Med Hypotheses. 2004;63(1):8–20. Epub 2004/06/15. doi: 10.1016/j.mehy.2003.12.045. PubMed PMID: 15193340.

Poole AC, Thomas RE, Andrews LA, McBride HM, Whitworth AJ, Pallanck LJ. The PINK1/Parkin pathway regulates mitochondrial morphology. Proc Natl Acad Sci U S A. 2008;105(5):1638–43. Epub 2008/01/31. doi: 10.1073/pnas.0709336105. PubMed PMID: 18230723; PMCID: PMC2234197.

Magrane J, Manfredi G. Mitochondrial function, morphology, and axonal transport in amyotrophic lateral sclerosis. Antioxid Redox Signal. 2009;11(7):1615–26. Epub 2009/04/07. doi: 10.1089/ARS.2009.2604. PubMed PMID: 19344253; PMCID: PMC2789440.

Ramonet D, Perier C, Recasens A, Dehay B, Bove J, Costa V, Scorrano L, Vila M. Optic atrophy 1 mediates mitochondria remodeling and dopaminergic neurodegeneration linked to complex I deficiency. Cell Death Differ. 2013;20(1):77–85. Epub 2012/08/04. doi: 10.1038/cdd.2012.95. PubMed PMID: 22858546; PMCID: PMC3524632.

Lane DJ, Huang ML, Ting S, Sivagurunathan S, Richardson DR. Biochemistry of cardiomyopathy in the mitochondrial disease Friedreich’s ataxia. Biochem J. 2013;453(3):321–36. Epub 2013/07/16. doi: 10.1042/BJ20130079. PubMed PMID: 23849057.

Von Stockum S, Nardin A, Schrepfer E, Ziviani E. Mitochondrial dynamics and mitophagy in Parkinson’s disease: A fly point of view. Neurobiol Dis. 2016;90:58–67. Epub 2015/11/10. doi: 10.1016/j.nbd.2015.11.002. PubMed PMID: 26550693.

Camilleri A, Ghio S, Caruana M, Weckbecker D, Schmidt F, Kamp F, Leonov A, Ryazanov S, Griesinger C, Giese A, Cauchi RJ, Vassallo N. Tau-induced mitochondrial membrane perturbation is dependent upon cardiolipin. Biochim Biophys Acta Biomembr. 2020;1862(2):183064. Epub 2019/09/16. doi: 10.1016/j.bbamem.2019.183064. PubMed PMID: 31521630.

Wertz MH, Mitchem MR, Pineda SS, Hachigian LJ, Lee H, Lau V, Powers A, Kulicke R, Madan GK, Colic M, Therrien M, Vernon A, Beja-Glasser VF, Hegde M, Gao F, Kellis M, Hart T, Doench JG, Heiman M. Genome-wide In Vivo CNS Screening Identifies Genes that Modify CNS Neuronal Survival and mHTT Toxicity. Neuron. 2020;106(1):76–89 e8. Epub 2020/02/01. doi: 10.1016/j.neuron.2020.01.004. PubMed PMID: 32004439; PMCID: PMC7181458.

Winquist RJ, Gribkoff VK. Targeting putative components of the mitochondrial permeability transition pore for novel therapeutics. Biochem Pharmacol. 2020;177:113995. Epub 2020/04/28. doi: 10.1016/j.bcp.2020.113995. PubMed PMID: 32339494.

Browne SE, Bowling AC, MacGarvey U, Baik MJ, Berger SC, Muqit MM, Bird ED, Beal MF. Oxidative damage and metabolic dysfunction in Huntington’s disease: selective vulnerability of the basal ganglia. Ann Neurol. 1997;41(5):646–53. Epub 1997/05/01. doi: 10.1002/ana.410410514. PubMed PMID: 9153527.

Mann VM, Cooper JM, Javoy-Agid F, Agid Y, Jenner P, Schapira AH. Mitochondrial function and parental sex effect in Huntington’s disease. Lancet. 1990;336(8717):749. Epub 1990/09/22. doi: 10.1016/0140-6736(90)92242-a. PubMed PMID: 1975918.

Brennan WA, Jr., Bird ED, Aprille JR. Regional mitochondrial respiratory activity in Huntington’s disease brain. J Neurochem. 1985;44(6):1948–50. Epub 1985/06/01. doi: 10.1111/j.1471-4159.1985.tb07192.x. PubMed PMID: 2985766.

Shirendeb U, Reddy AP, Manczak M, Calkins MJ, Mao P, Tagle DA, Reddy PH. Abnormal mitochondrial dynamics, mitochondrial loss and mutant huntingtin oligomers in Huntington’s disease: implications for selective neuronal damage. Hum Mol Genet. 2011;20(7):1438–55. Epub 2011/01/25. doi: 10.1093/hmg/ddr024. PubMed PMID: 21257639; PMCID: PMC3049363.

Agrawal S, Fox J, Thyagarajan B, Fox JH. Brain mitochondrial iron accumulates in Huntington’s disease, mediates mitochondrial dysfunction, and can be removed pharmacologically. Free Radic Biol Med. 2018;120:317–29. Epub 2018/04/07. doi: 10.1016/j.freeradbiomed.2018.04.002. PubMed PMID: 29625173; PMCID: PMC5940499.

Browne SE. Mitochondria and Huntington’s disease pathogenesis: insight from genetic and chemical models. Ann N Y Acad Sci. 2008;1147:358–82. Epub 2008/12/17. doi: 10.1196/annals.1427.018. PubMed PMID: 19076457.

Zarate N, Gomez-Pastor R. Excitatory synapse impairment and mitochondrial dysfunction in Huntington’s disease: heat shock factor 1 (HSF1) converging mechanisms. Neural Regen Res. 2020;15(1):69–70. Epub 2019/09/20. doi: 10.4103/1673-5374.264459. PubMed PMID: 31535651; PMCID: PMC6862424.

Franco-Iborra S, Plaza-Zabala A, Montpeyo M, Sebastian D, Vila M, Martinez-Vicente M. Mutant HTT (huntingtin) impairs mitophagy in a cellular model of Huntington disease. Autophagy. 2020:1–18. Epub 2020/02/26. doi: 10.1080/15548627.2020.1728096. PubMed PMID: 32093570.

Agrawal S, Fox JH. Novel proteomic changes in brain mitochondria provide insights into mitochondrial dysfunction in mouse models of Huntington’s disease. Mitochondrion. 2019;47:318–29. Epub 2019/03/25. doi: 10.1016/j.mito.2019.03.004. PubMed PMID: 30902619; PMCID: PMC7011782.

Koc EC, Koc H. Regulation of mammalian mitochondrial translation by post-translational modifications. Biochim Biophys Acta. 2012;1819(9-10):1055–66. Epub 2012/04/07. doi: 10.1016/j.bbagrm.2012.03.003. PubMed PMID: 22480953.

Suen DF, Norris KL, Youle RJ. Mitochondrial dynamics and apoptosis. Genes Dev. 2008;22(12):1577–90. Epub 2008/06/19. doi: 10.1101/gad.1658508. PubMed PMID: 18559474; PMCID: PMC2732420.

Hamilton J, Brustovetsky T, Khanna R, Brustovetsky N. Mutant huntingtin does not cross the mitochondrial outer membrane. Hum Mol Genet. 2020;29(17):2962–75. Epub 2020/08/22. doi: 10.1093/hmg/ddaa185. PubMed PMID: 32821928; PMCID: PMC7566381.

Yu ZX, Li SH, Evans J, Pillarisetti A, Li H, Li XJ. Mutant huntingtin causes context-dependent neurodegeneration in mice with Huntington’s disease. J Neurosci. 2003;23(6):2193–202. Epub 2003/03/27. PubMed PMID: 12657678; PMCID: PMC6742008.

Couvillion MT, Soto IC, Shipkovenska G, Churchman LS. Synchronized mitochondrial and cytosolic translation programs. Nature. 2016;533(7604):499–503. Epub 2016/05/27. doi: 10.1038/nature18015. PubMed PMID: 27225121; PMCID: PMC4964289.

Ostojic J, Panozzo C, Bourand-Plantefol A, Herbert CJ, Dujardin G, Bonnefoy N. Ribosome recycling defects modify the balance between the synthesis and assembly of specific subunits of the oxidative phosphorylation complexes in yeast mitochondria. Nucleic Acids Res. 2016;44(12):5785–97. Epub 2016/06/04. doi: 10.1093/nar/gkw490. PubMed PMID: 27257059; PMCID: PMC4937339.

Desai N, Yang H, Chandrasekaran V, Kazi R, Minczuk M, Ramakrishnan V. Elongational stalling activates mitoribosome-associated quality control. Science. 2020;370(6520):1105–10. Epub 2020/11/28. doi: 10.1126/science.abc7782. PubMed PMID: 33243891; PMCID: PMC7116630.

Trettel F, Rigamonti D, Hilditch-Maguire P, Wheeler VC, Sharp AH, Persichetti F, Cattaneo E, MacDonald ME. Dominant phenotypes produced by the HD mutation in STHdh(Q111) striatal cells. Hum Mol Genet. 2000;9(19):2799–809. Epub 2000/11/25. doi: 10.1093/hmg/9.19.2799. PubMed PMID: 11092756.

Pryor WM, Biagioli M, Shahani N, Swarnkar S, Huang WC, Page DT, MacDonald ME, Subramaniam S. Huntingtin promotes mTORC1 signaling in the pathogenesis of Huntington’s disease. Sci Signal. 2014;7(349):ra103. Epub 2014/10/30. doi: 10.1126/scisignal.2005633. PubMed PMID: 25351248.

Guo H, Ingolia NT, Weissman JS, Bartel DP. Mammalian microRNAs predominantly act to decrease target mRNA levels. Nature. 2010;466(7308):835–40. Epub 2010/08/13. doi: 10.1038/nature09267. PubMed PMID: 20703300; PMCID: PMC2990499.

Miettinen TP, Bjorklund M. Modified ribosome profiling reveals high abundance of ribosome protected mRNA fragments derived from 3’ untranslated regions. Nucleic Acids Res. 2015;43(2):1019–34. Epub 2015/01/01. doi: 10.1093/nar/gku1310. PubMed PMID: 25550424; PMCID: PMC4333376.

Martin M. Cutadapt removes adapter sequences from high-throughput sequencing reads. EMBnetjournal. 2011;17(1):10–2.

Li B, Dewey CN. RSEM: accurate transcript quantification from RNA-Seq data with or without a reference genome. BMC Bioinformatics. 2011;12:323. Epub 2011/08/06. doi: 10.1186/1471-2105-12-323. PubMed PMID: 21816040; PMCID: PMC3163565.

Quinlan AR, Hall IM. BEDTools: a flexible suite of utilities for comparing genomic features. Bioinformatics. 2010;26(6):841–2. Epub 2010/01/30. doi: 10.1093/bioinformatics/btq033. PubMed PMID: 20110278; PMCID: PMC2832824.

Love MI, Huber W, Anders S. Moderated estimation of fold change and dispersion for RNA-seq data with DESeq2. Genome Biol. 2014;15(12):550. Epub 2014/12/18. doi: 10.1186/s13059-014-0550-8. PubMed PMID: 25516281; PMCID: PMC4302049.

